# *Communication*: Theoretical Insights from Binding of *Neigleria fowleri* Cell Division Proteins with Amoebicidal Quassinoids

**DOI:** 10.1101/367912

**Authors:** Zarrin Basharat, Shumaila Zaib, Azra Yasmin

**Affiliations:** Jamil-ur-Rahman center for Genome Research, Dr. Panjwani Centre for Molecular Medicine and Drug Research, International Center for Chemical and Biological Sciences, University of Karachi-75270, Karachi, Pakistan.; Microbiology & Biotechnology Research Lab, Department of Environmental Sciences, Fatima Jinnah Women University, Rawalpindi 46000, Pakistan.

**Keywords:** *Neigleria fowleri*, Brain eating amoeba, Primary amoebic meningoencephalitis, Quassinoids, Amoebicidal, Phytotherapy, Molecular docking

## Abstract

The ameboflagellate *Neigleria fowleri*, also known as brain eating amoeba is responsible for fatal primary amoebic meningoencephalitis (PAM) infection in humans. Cell division proteins (CDPs) in *N. fowleri* have been uncharacterized until now, despite their importance in the initiation of cell division and proliferation of the pathogen. Here, we report characterization and structural assembly of eight such proteins associated with division and docking them with anti-amoebic quassinoid compounds. Quassinoids have been implicated as inhibitors of cell proliferation of amoeboid species as well as tumor cells. Here, they were screened computationally to find interaction mechanism as well as binding energies with CDPs of *N. fowleri*. The identified inhibitors could play a role in prevention of cell division and hence, stop *N. fowleri* growth and proliferation during infection. This study supports CDPs as a target for anti-amoebic intervention and identifies quassinoid phytochemical compounds as suitable for optimization into a new therapy against *N. fowleri*.

## 1. Introduction

*N. fowleri* is considered as an amphizoic amoeba because of its property to survive in a free-living state as well as in the host. In free living form it is found in water or soil and in the host found in human central nervous system [1]. Life cycle of *N. fowleri* is comprised of three stages, Trophozoites (feeding stage), Flagellate (motility stage) and Cysts (surviving). *N. fowleri* enters the human body in the trophozoite form which is infectious stage. Food cups are found on the surface of trophozoites through which they ingest bacteria, fungi, and human tissue [2]. *N. fowleri* passes into the host during recreational activities such as water sports, swimming, cultural practices (use of neti pots) religious practices of Muslims (ablution or “wuzu”) and Hindus (bathing in river Ganges) [3]. *N. fowleri* forces itself into the nasal cavity and after passing through olfactory nerve *N. fowleri* reaches CNS via cribriform plate [2]. This pathogen attacks by releasing cytotoxic molecules in the central nervous system which cause necrosis and lead to primary amoebic meningoencephalitis [4].

PAM was reported for first time in Australia in 1965 although there has been a record of its occurrance as early as 1937 in Virginia, America. Cases have been reported from the America, Australia, Europe and Asia. Rise in temperature is the major factor which contributes to increases PAM infection, as high temperature favours multiplication of the amoeba [5,6]. PAM symptoms are fever, nausea, vomiting, severe headache, stiff neck, altered mental status, hallucinations, seizures, and coma [7]. Overall, this disease is extremely rare but deadly; about 235 PAM cases have been reported worldwide till to date. Reports have been of almost always fatal PAM infection, with survival rate of approximately 5% [8]. In Pakistan alone, 40 deaths have been reported in Karachi from 2010 to 2015 [7]. The incubation period for PAM is about 5 to 7 days, and infected individual died within a week [1]. From 1962 to 2014, only three people out of 133 infected with *N. fowleri* have survived in the USA [6]. Cerebrospinal fluid smear, PCR and CT scan are normally used to diagnose *N. fowleri* ([7].

Drugs that have been used in the past for treatment of PAM inlcude Amphotericin B, Fluconazole, Azithromycin, Miltefosine and Rifamin. These drugs either alter membrane permeability, inhibit protein synthesis or RNA polymerase activity, with side effects like diarrhoea, nausea, chills, vomiting, malaise, heartburn, drowsiness, dizziness and headache [4]. Owing to lack of any vaccine and side effects of antibiotics against *N. fowleri* [9], improved drugs are required for prevention and treatment of *N. fowleri* infection. For the discovery and development of drugs against selected targets, plethora of computational approaches such as comparative genomics, molecular docking and virtual screening are used. Previously, we reported HSP70 as potential therapeutic candidate against *N. fowleri* [10]. CDPs of *N. fowleri* could also serve as important therapeutic candidates. In this study, anti-amoebic quassinoids from literature were mined and used for molecular docking with CDPs of *N. fowleri*. Currently, the authors are also involved in screening large library of quassinoids against different amoebic species and their comparison at ortholog and paralog scale.

## 2. Method and analysis

Interest in computational drug discovery has been increasing in the recent era and chemicals scaffolds are being explored by researchers for their therapeutic and pharmacological effects [11] using various computational algorithms and softwares. For computational assisted screening of quassinoids against CDPs of *N. fowleri*, CDP sequences were downloaded from the Amoeba Genomics Resource Database (http://amoebadb.org/amoeba/). Accession numbers have been mentioned in the Table 1. Phsphorylation potential was obtained from the NetPhos 3.1 server [12]. Interplay of phosphorylation and O-β-glycosylation i.e. Yin-Yang sites were identified using YinOYang server 2.1 [13]. Molecular weight, theoretical pI and grand average of hydrophobicity (GRAVY) were identified using Expasy tools [14]. TMHMM server [15] was used for assessing the presence of transmembrane helices in the proteins.

3D structures were predicted using the I-TASSER software suit [16, 17] using templates from the RCSB Protein Data Bank. Quassinoid compounds (Table 2) were downloaded from the PubChem database, optimized and stored in a .mdb database. For docking, protein structures were protonated as well as energy minimized until a gradient of 0.05 was attained. Docking was carried out in Molecular Operating Environment (MOE) software suite. Parameters for placement: Triangle match and rescoring by London dG and Affinity dG force respectively. Refinement was done through forcefield method. The best structure with least S value was retained for further downstream analysis. Ligand interactions were visualized in 2D [18].

**Table 1.** Different physico-chemical parameters of *N. fowleri* CDPs. Residues in brackets against the third and sixth CDP are the ones showing phosphorylation at residues interacting with quassinoid molecules.

| Serial No. | Accession no. | No. of amino acids | Molecular weight | Theoretical pI | GRAVY | Predicted TMHs | No. of phosphorylated residues | No. of YinOYang sites |
| --- | --- | --- | --- | --- | --- | --- | --- | --- |
| 1 | NF0026440 | 545 | 60032.69 | 5.23 | -0.579 | 0 | 106 | 25 |
| 2 | NF0036940 | 382 | 45053.14 | 8.64 | -0.188 | 3 | 39 | 3 |
| 3 | NF0045650 | 161 | 19343.08 | 5.60 | -0.635 | 0 | 21 (T29, T30) | 3 |
| 4 | NF0080760_2 | 258 | 29006.76 | 6.54 | -0.301 | 0 | 31 | 6 |
| 5 | NF0084440_2 | 137 | 15949.08 | 8.90 | 0.803 | 2 | 8 | - |
| 6 | NF0090830 | 398 | 43473.21 | 9.89 | -0.892 | 0 | 79 (T341, S346) | 18 |
| 7 | NF0111200 | 384 | 42852.93 | 7.10 | -0.439 | 0 | 40 | 2 |
| 8 | NF0120690_1 | 106 | 12601.94 | 8.58 | 0.460 | 2 | 8 | - |

**Table 2.**
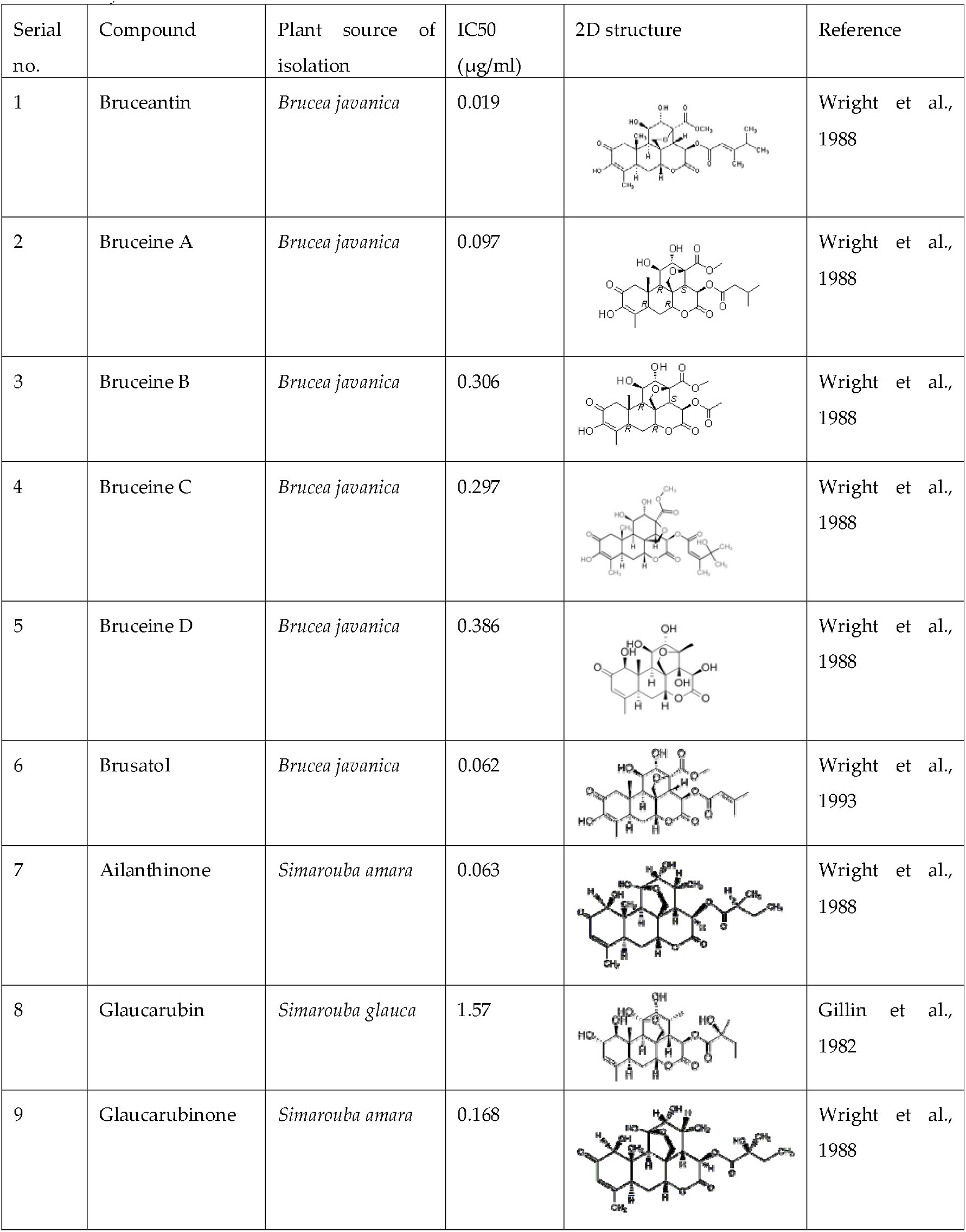
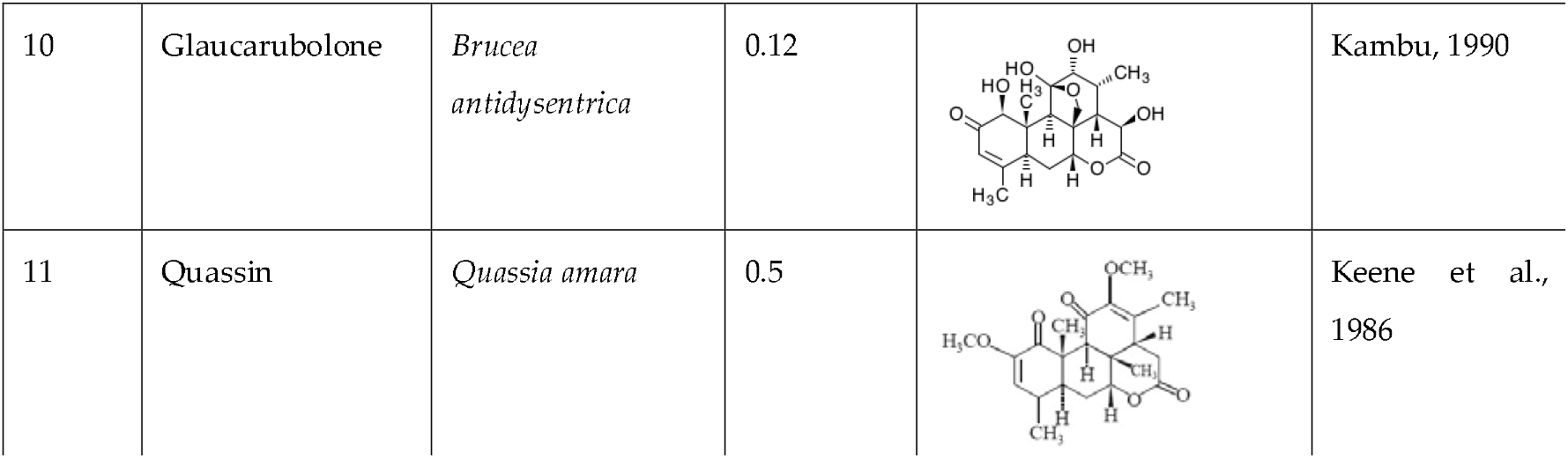
Anti-amoebic quassinoids (Reviewed by [19]), used in this study for docking and analysis against CDPs of *N. fowleri*.

| Serial no. | Compound | Plant source of isolation | IC50 (µg/ml) | 2D structure | Reference |
| --- | --- | --- | --- | --- | --- |
| 1 | Bruceantin | *Brucea javanica* | 0.019 |  | Wright et al., 1988 |
| 2 | Bruceine A | *Brucea javanica* | 0.097 |  | Wright et al., 1988 |
| 3 | Bruceine B | *Brucea javanica* | 0.306 |  | Wright et al., 1988 |
| 4 | Bruceine C | *Brucea javanica* | 0.297 |  | Wright et al., 1988 |
| 5 | Bruceine D | *Brucea javanica* | 0.386 |  | Wright et al., 1988 |
| 6 | Brusatol | *Brucea javanica* | 0.062 |  | Wright et al., 1993 |
| 7 | Ailanthinone | *Simarouba amara* | 0.063 |  | Wright et al., 1988 |
| 8 | Glaucarubin | *Simarouba glauca* | 1.57 |  | Gillin et al., 1982 |
| 9 | Glaucarubinone | *Simarouba amara* | 0.168 |  | Wright et al., 1988 |
| 10 | Glaucarubolone | *Brucea antidysentrica* | 0.12 |  | Kambu, 1990 |
| 11 | Quassin | *Quassia amara* | 0.5 |  | Keene et al., 1986 |

## 3. Results

We are constantly in search of new and better drugs for cure against diseases of almost all types. High-throughput screening, structure based interaction and quantitative structure activity relationship is usually used for guiding bioassays. Isolated molecules may then be developed into drugs with improved clinical action or fewer side effects [19]. There is a lot of scope in investigating the nature derived plant compounds and herbal therapies for their safety and efficacy and harnessing their potential against diseases. Several drugs have been tested for *N. fowleri* caused PAM with varying results [20–30].

Quassinoids are phytoconstituents of several Simaroubaceae species and consist of degraded triterpenes. They have demonstrated activity against viral proteins [31, 32] and cancerous cells [33,34]. They have also shown pharmacological action against inflammation, insects and ulcer [31]. They bind proteins in amoeba and prevent their proliferation. Their impact on protozoan/amoeba has been reviewed in detail elsewhere [19]. In the quest for new, effective, and safe amoebicidal agents against *N. fowleri*, anti-amoebic quassinoid molecule interactions were identified against CDPs of this pathogen. Each CDP was tested against all eight anti-ameobic quassinoids (mentioned in Table 2) and small molecule with the best ability to bind to the specific CDP was analysed for interacting residues.

All CDPs had different sequence and hence, structures. This demonstrates a pattern of paralog existence and extreme variation of CDP sequences in *N. fowleri*. Molecular interaction did not show conservation of a specific binding residue. Despite this variation of sequence and no conservation of binding site, majority i.e. 3 paralogs showed best affinity for Brucein B molecule. Two CDPs showed maximum affinity for Brusatol (Table 3). Phosphorylation pattern also varied for all paralogs. In two CDPs (Accession no: NF0045650 and NF0090830), two phosphorylated residues made interaction with ligand. One had both threonines while other paralog had one serine and one threonine showing interaction with quassinoids Glaucarubin and Bruceine B respectively. No such pattern was observed for other paralogs. YinOYang sites were present in six CDPs (Table 3) but none showed interplay of phosphorylation and glycosylation at docked site residues.

**Table 3.**
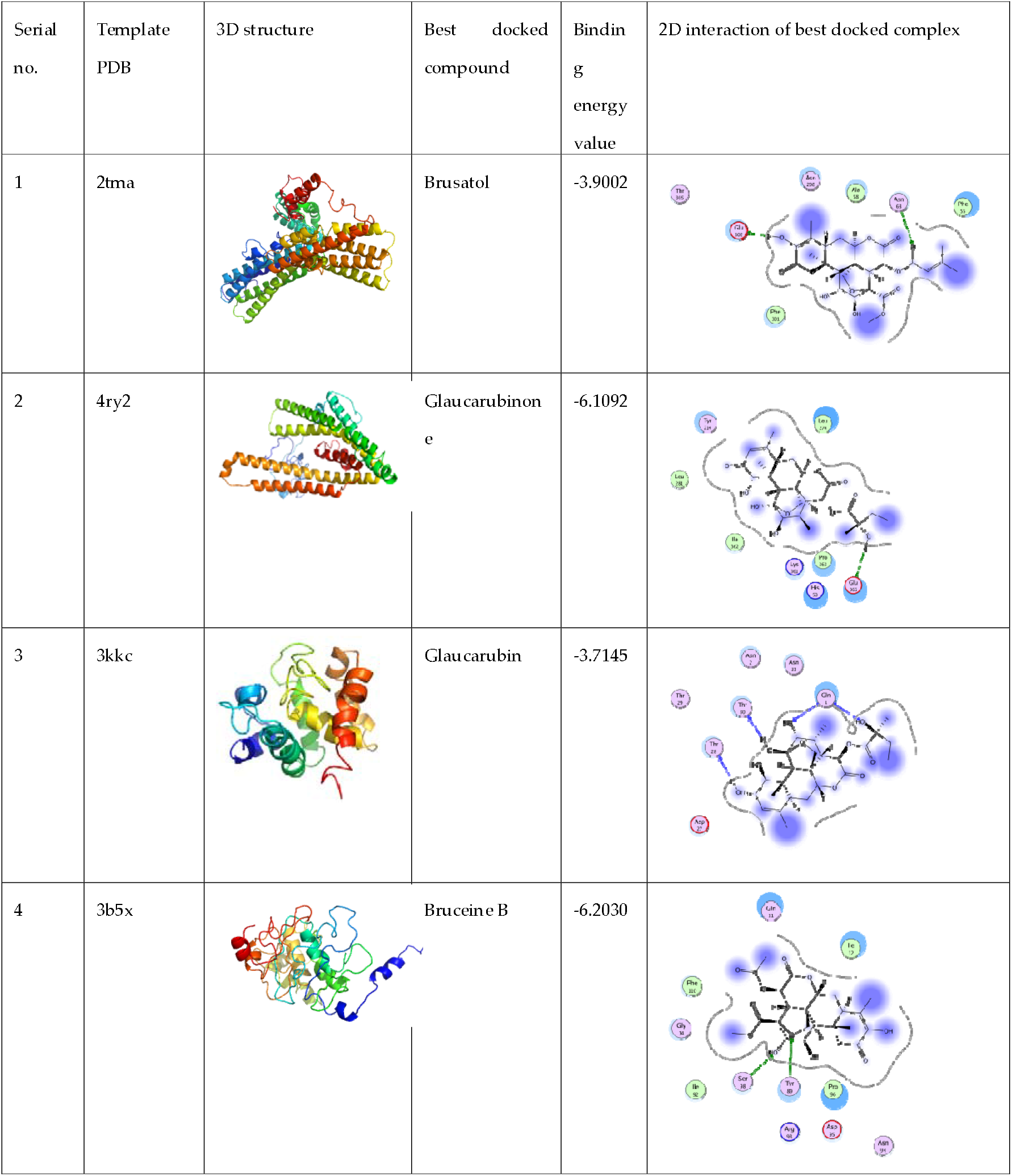
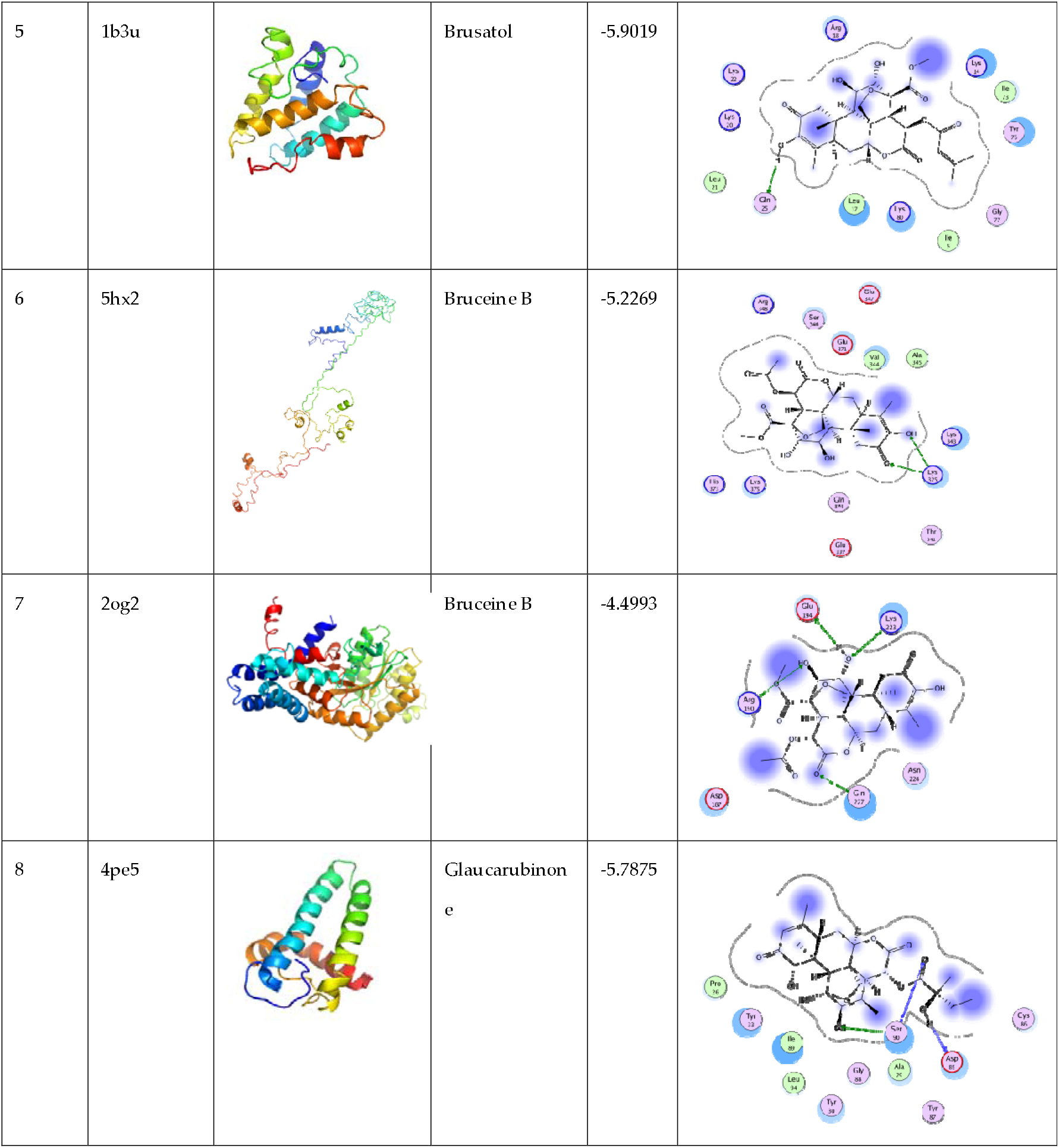
Protein structure and interaction information for the studied CDPs of *N. fowleri*. Serial numbers refer to the accession numbers mentioned against similar serial numbers in Table 1. 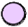 depicts polar residue, 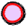 acidic, 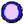 basic, 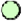 hydrophobic, 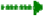 side chain acceptor, 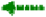 side chain donor, 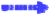 backbone acceptor, 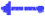 backbone donor, 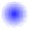 ligand exposure, 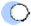 receptor exposure, 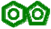 arene-arene, 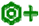 arene cation and black dotted line as proximity contour.

Since, formulation of plant derived small molecules into drugs, entails screening enormous number of plant extracts followed by the compound isolation and identification of action mechanism, this study could prove useful for guiding such assays. Toxicity tests could then be done to establish safety for the host and specificity of action on pathogen. In the present study, docking was attempted to screen potent amoebicidal quassinoid phytochemicals against CDPs of *N. fowleri*. The study is a preliminary information report and experimental assays are further needed to validate the findings.

## Supplementary Materials

The predicted 3D protein structure coordinates of cell division proteins (Serial no. 1–8, Table 1) can be requested from the corresponding author.

## Acknowledgments

The authors are thankful to Alexandra Elbakyan for literature support. This work did not receive any particular funding.

## Author Contributions

“Z.B. and A.Y. conceived and designed the experiments; Z.B. performed the experiments; Z.B. analyzed the data; Z.B. and S.Z. wrote the original draft of the paper; AY edited and checked the final version of paper.”

